# SARS-CoV-2-associated ssRNAs activate inflammation and immunity via TLR7/8

**DOI:** 10.1101/2021.04.15.439839

**Authors:** Valentina Salvi, Hoang Oanh Nguyen, Francesca Sozio, Tiziana Schioppa, Mattia Laffranchi, Patrizia Scapini, Mauro Passari, Ilaria Barbazza, Laura Tiberio, Nicola Tamassia, Cecilia Garlanda, Annalisa Del Prete, Marco A. Cassatella, Alberto Mantovani, Silvano Sozzani, Daniela Bosisio

## Abstract

The inflammatory and IFN pathways of innate immunity play a key role in both resistance and pathogenesis of Coronavirus Disease 2019 (COVID-19). Innate sensors and SARS-CoV-2-Associated Molecular Patterns (SAMPs) remain to be completely defined. Here we identify single-stranded RNA (ssRNA) fragments from SARS-CoV-2 genome as direct activators of endosomal TLR7/8 and MyD88 pathway. The same sequences induced human DC activation in terms of phenotype and functions, such as IFN and cytokine production and Th1 polarization. A bioinformatic scan of the viral genome identified several hundreds of fragments potentially activating TLR7/8, suggesting that products of virus endosomal processing potently activate the IFN and inflammatory responses downstream these receptors. In vivo, SAMPs induced MyD88-dependent lung inflammation characterized by accumulation of proinflammatory and cytotoxic mediators and immune cell infiltration, as well as splenic DC phenotypical maturation. These results identify TLR7/8 as crucial cellular sensors of ssRNAs encoded by SARS-CoV-2 involved in host resistance and disease pathogenesis of COVID-19.

## Introduction

SARS coronavirus 2 (SARS-CoV-2), is a positive-sense ssRNA virus belonging to the family of Coronaviridae, also including the closely related Middle East respiratory syndrome coronavirus (MERS-CoV) and SARS-CoV **(1)**. In a subgroup of patients, SARS-CoV-2 infection (Coronavirus disease 2019, COVID-19) develops as acute respiratory distress syndrome featuring intense lung injury, sepsis-like manifestations and multi-organ failure (2) associated with overt production of pro-inflammatory cytokines that directly correlates with poor prognosis (3). This clinical condition suggests that an overactive innate immune response may unleash virus-dependent immune pathology (4). Innate immune activation is also responsible for inducing the protective antiviral state, largely mediated by the release of type I IFNs. Indeed, inborn errors in type I IFN production and amplification (5) or pre-existing blocking auto-antibodies against members of the IFN family of cytokines (6) were found to correlate with unfavorable prognosis.

DCs act as crucial messengers linking innate and adaptative immunity against viral infections (7, 8). Within DC heterogeneity, plasmacytoid DCs (pDCs) play an important role as the major source of type I IFN in response to viral infection, while conventional DCs (cDCs) respond to a vast variety of pathogens by producing pro-inflammatory cytokines and are the main responsible for T cell activation (9–11). pDCs sense ssRNA viruses through TLR7 (12), an endosomal receptor activated by genomic fragments rich in guanine (G) and uracil (U), derived by endosomal processing of the virus independently of infection (13). By contrast, cDCs express the closely related TLR8 (14). Despite the fact that TLR7 and TLR8 display high structural and functional homology, similar ligand specificity (15) and recruit the same signaling intracellular adaptor molecule, MyD88 (16), the signaling pathways of these two TLRs diverge in the functional significance, with TLR7 more involved in the antiviral immune response and TLR8 mastering the production of pro-inflammatory cytokines. Both cDCs and pDCs were shown to be reduced in the blood of severe acute COVID-19 patients (17, 18) as a possible result of cell activation (19), but the mechanisms of SARS-CoV-2 recognition and activation by innate immune cells still need to be identified. This study characterizes the first SARS-CoV-2-associated molecular patterns (SAMPs) and identifies the TLR7/8/MyD88 axis as a crucial pathway in the activation of human pDCs and cDCs.

## Results

### Identification of potential ssRNA SAMPs

Based on previous work identifying RNA40, a ssRNA rich in guanine and uracil (GU-rich) from the U5 region of HIV-1, as the first natural agonist of TLR7 and TLR8 (20) and on known features of TLR7/8 ligands (15, 21, 22), we searched for putative immunostimulatory sequences within the SARS-CoV-2 ssRNA genome. Our bioinformatic scan revealed 491 GU-rich sequences, among which more than 250 also bearing at least one “UGUGU” Interferon Induction Motif (IIM) (15, 20, 21) (Suppl. Table 1).

We hypothesized that these sequences may represent so far unidentified SAMPs responsible for viral recognition and immune activation via endosomal TLR triggering. The elevated number of sequences detected suggests that, upon endosomal engulfment, the fragmentation of the SARS-CoV-2 genome may generate many TLR7/8-triggering sequences, thus displaying high chances to contact and activate the IFN and inflammatory responses downstream these receptors.

To validate the stimulatory potential on innate immune cells, two representative sequences, SCV2-RNA1 and SCV2-RNA2, were chosen within the previous list, synthesized and tested in *in vitro* and *in vivo* models of inflammation.

### ssRNA SAMPs activate human monocyte-derived DCs (moDCs)

moDCs, a model of inflammatory cDCs expressing a wide variety of TLRs (7, 23–25), were treated with increasing concentrations of SCV2-RNA1 and SCV2-RNA2 along with HIV-1-derived RNA40 (Heil 2004), used as a positive control. U/A alternated control sequences SCV2-RNA1A and SCV2-RNA2A were used as negative controls (see materials and methods). Figure 1A shows that both fragments efficiently activated cytokine secretion by moDCs. In particular, we observed potent induction of pro-inflammatory cytokines (TNF-α, IL-6), of the Th1-polarizing cytokine IL-12 and chemokines recruiting polymorphonuclear neutrophils (CXCL8), myelomonocytic cells (CCL3) and Th1- and cytotoxic effectors cells (CXCL9). Especially at low concentrations, SCV2-RNA1 and SCV2-RNA2 were more efficient than HIV-1-derived RNA40. In all experimental conditions, U/A alternated SCV2-RNA1A and SCV2-RNA2A did not induce cytokine secretion. SCV2-RNA1 and SCV2-RNA2 also induced moDC phenotypical maturation in terms of CD83, CD86 and CCR7 expression (Figure 1B). Similarly to cytokine secretion, upregulation of maturation markers by RNA40 was less effective. These results demonstrated that both SCV2-RNA1 and SCV2-RNA2 behave as SAMPs endowed with potent DC stimulatory capacity. Because of their similar potency, further experiments were carried out using a mixture of the two SAMPs (indicated as SCV2-RNA), a condition that may also better mimic a physiological stimulation by multiple sequences derived from SARS-CoV-2 genome endosomal fragmentation.

**Figure 1.**
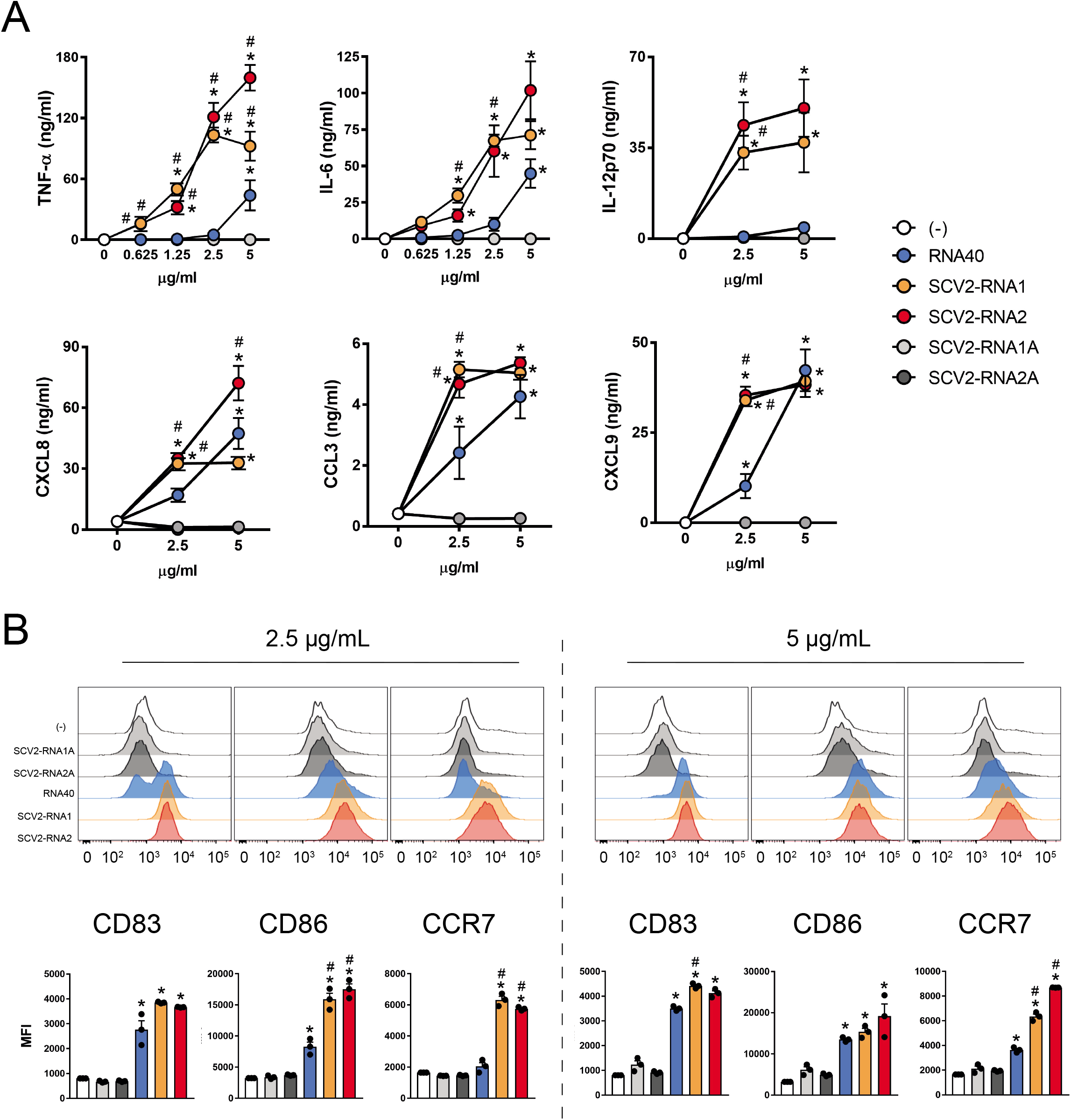
SAMPs activate cytokine secretion and phenotypical maturation of moDCs. (A) moDCs (2×10^6^/ml) were stimulated with increasing concentrations of the indicated viral RNAs or with vehicle alone (-) for 24 hours. The production of TNF-α, IL-6, IL-12p70, CXCL8, CCL3 and CXCL9 was evaluated by ELISA in cell-free supernatants. Data are expressed as mean ± SEM (n=3). Results of SCV2-RNA1A and SCV2-RNA2A are superimposed in all graphs. (B) moDCs were stimulated as described in (A) and the surface expression of CD83, CD86 and CCR7 evaluated by FACS analysis. Data are expressed as representative cytofluorimetric profiles (upper panels) or as the mean ± SEM (n=3) of the Median of Fluorescence Intensity (MFI) (lower panels). (A-B) *P< 0.05 versus (-) by one-way ANOVA with Dunnett’s post-hoc test; #P< 0.05 versus RNA40 by paired Student’s *t* test.

### ssRNA SAMPs activate T cell responses

The impact of SAMPs on the ability of DCs to stimulate T cell functions was investigated in co-culture experiments of SAMP-activated DCs with allogeneic naïve CD4^+^ and CD8^+^ T cells. Figure 2A shows that SAMP-activated DCs induced proliferation of both naïve CD4^+^ (left) and CD8^+^ (right) T cells. Activated CD4^+^ T cells produced IFN-γ but no IL-4, a typical Th1-effector phenotype (Figure 2B). Functional activation of CD8^+^ T cells was similarly demonstrated by the detection of secreted IFN-γ (Figure 2C, left panel) and the intracellular accumulation of Granzyme B (GrB, right panel), a marker of a cytotoxic phenotype. None of these effects were observed when DCs were activated with U/A alternated SAMPs.

**Figure 2.**
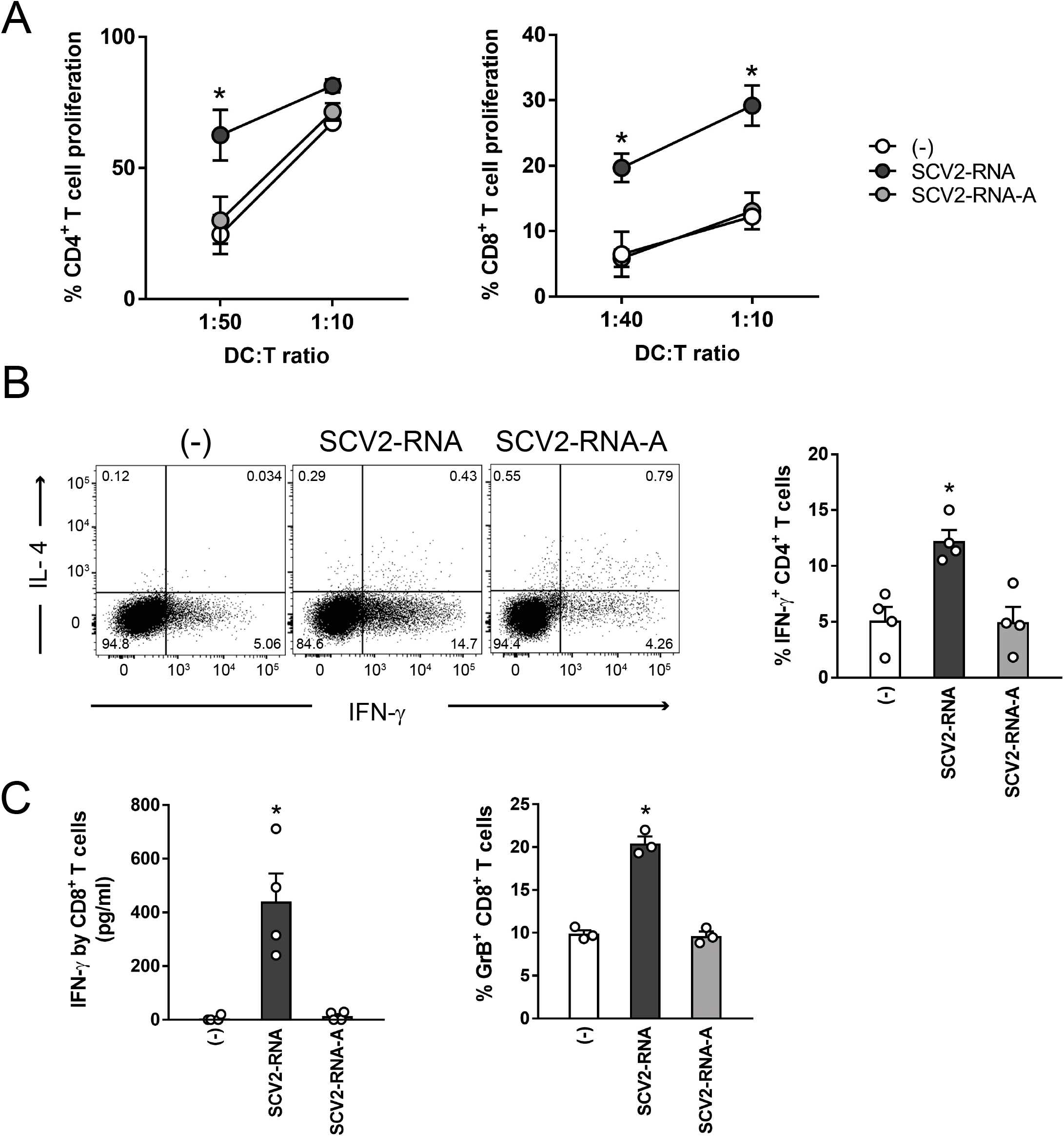
SAMP-activated DCs trigger T cell proliferation and functional activation. (A) moDCs were stimulated with vehicle (-) or with SCV2-RNA or the A-to-U-replaced SCV2-RNA-A (both at 5 μg/ml) for 24 hours. Activated moDCs were co-cultured for 6 days with CFSE-stained allogenic naïve CD4^+^T cells or CD8^+^ T cells at the indicated DC:T cell ratio. Alloreactive T cell proliferation was assessed by measuring CellTrace-CFSE dye loss by flow cytometry. Data are expressed as mean ± SEM (n=3) of the percentage of proliferating T cells. (B) moDCs stimulated as in (A) were cocultured for 6 days with allogenic naïve CD4^+^ T (DC:T cell ratio 1:20). Intracellular IFN-γ and IL-4 were evaluated by FACS analysis. Left, dot plots from one representative experiment. Right, bar graphs from four independent experiments. Data are expressed as mean ± SEM of the percentage of IFN-γ-producing cells. (C) moDCs activated as in (A) were cocultured for 6 days with allogenic CD8^+^ T (DC:T cell ratio 1:10). IFN-γ production was evaluated by ELISA in cell-free supernatants and intracellular Granzyme B (GrB) by FACS analysis. Data are expressed as mean ± SEM (n=3). (A-C) *P< 0.05 versus (-) by one-way ANOVA with Dunnett’s post-hoc test.

These experiments demonstrated that phenotypical DC maturation induced by SAMPs (Figure 1B) is paralleled by the acquisition of T-cell activating capabilities. Thus, SAMPs have the ability to induce a Th1-oriented immune response.

### ssRNA SAMPs activate human primary DCs

The ability of SCV2-RNAs to activate DCs was further investigated using primary circulating cDCs (comprising CD141^+^ cDC1 and CD1c^+^ cDC2) and BDCA2^+^ pDCs. SCV2-RNA efficiently induced the secretion of TNF-α and IL-6 (Figure 3A) and the expression of maturation markers, such as CD86 and CCR7 (Figure 3B) in cDCs. Similarly, SAMPs stimulated the release of IFN-α and TNF-α by pDCs (Figure 3C), as well as their maturation in terms of CD86 upregulation and BDCA2 reduction (Figure 3D). Similarly to previous results, U/A alternated control sequences did not activate cytokine production or maturation in either pDCs and cDCs (not shown).

**Figure 3.**
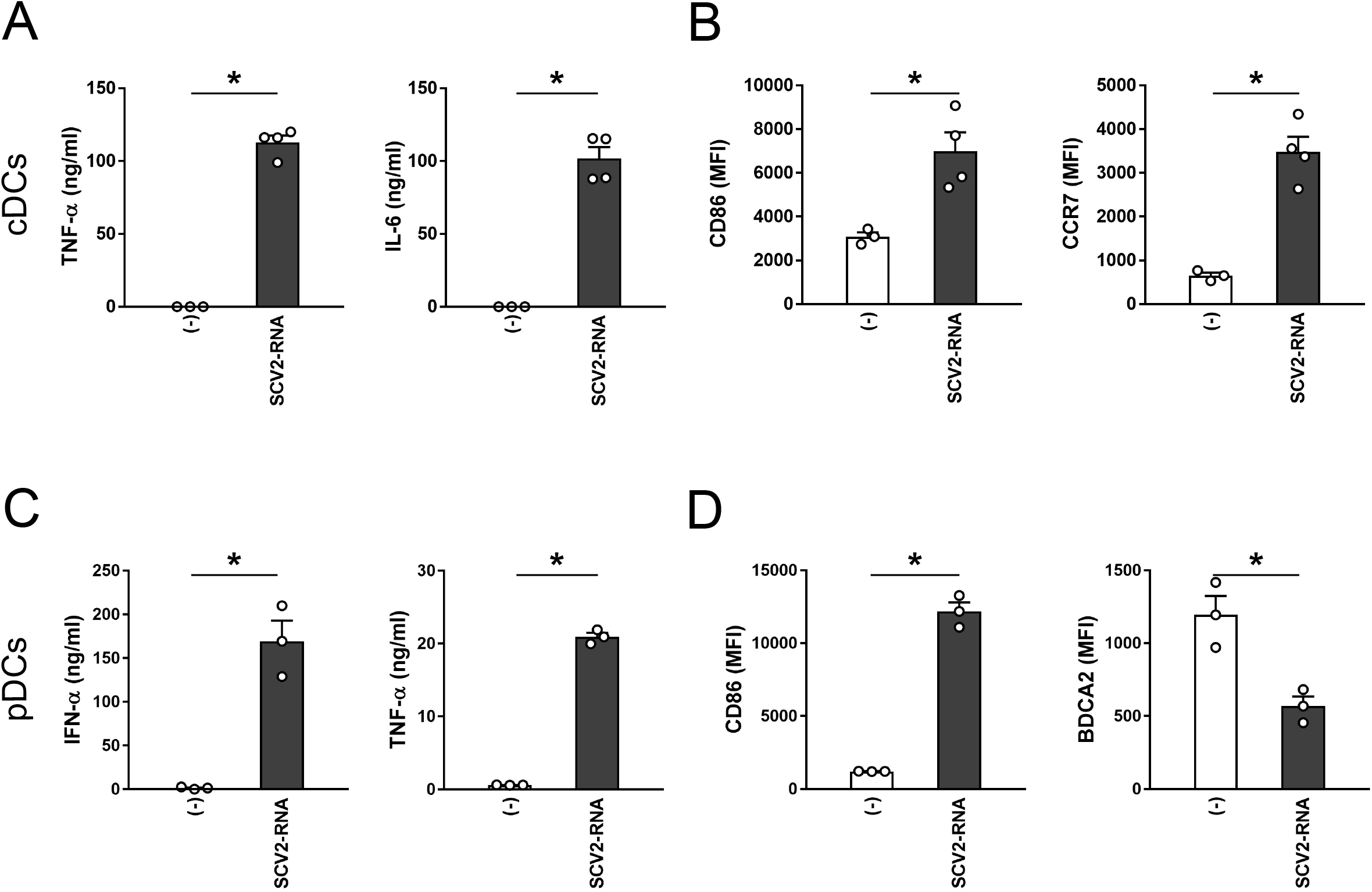
SAMPs activate cytokine secretion and phenotypical maturation in primary circulating DC subsets. cDCs (2×10^6^/ml) and pDCs (1×10^6^/ml) were stimulated with 5 μg/ml SCV2-RNA for 24 hours. (A-C) Cytokine secretion was evaluated by ELISA. Data are expressed as mean ± SEM (n=3-4); *P< 0.05 versus (-) by paired Student’s *t* test. (B-D) Surface expression of CD86, CCR7 and BDCA2 was evaluated by FACS analysis. Data are expressed as mean ± SEM (n=3-4) of the median fluorescence intensity (MFI); *P< 0.05 versus (-) by paired Student’s *t* test.

### ssRNA SAMPs act as TLR7/8 ligands

Despite the fact that SARS-CoV-2 has been shown to activate immune cells, the cellular sensors responsible for viral detection are still ill defined. To formally demonstrate the ability of SAMPs to functionally activate TLRs, experiments were performed in HEK-293 cells stably transfected with human TLR7 and TLR8 together with a NF-κB reporter gene. Figure 4A depicts the activation of NF-kB and luciferase production in both TLR7- and TLR8-expressing cells by SAMPs. Since both TLR7 and TLR8 signal through the common adaptor MyD88, siRNA interference was performed in moDCs. Figure 4B (left panel) shows that two different MyD88-specific siRNAs could decrease by about 50% the levels of MyD88 mRNA, while the expression of the TLR3-related adaptor TRIF was not affected. Consistent with this result, IL-6 production by SCV2-RNA was also decreased, whereas the stimulation of moDCs by Poly I:C was not affected (Figure 4B right panel). Next, moDCs were stimulated in the presence of CU-CPT9a, a specific TLR8 inhibitor (26). CU-CPT9a inhibited in a potent and dose-dependent manner the release of IL-6 when cells were stimulated with SCV2-RNA or R848 (TLR7/8 ligand). On the other hand, the TLR8 inhibitor did not affect the stimulation by LPS, a TLR4 ligand (Figure 4C). The complete inhibition observed upon TLR8 blocking is not surprising given the low expression of TLR7 in moDCs (23). TLR7 and TLR8 display a mutual exclusive expression in primary DCs. Indeed, cDCs express TLR8 as their unique endosomal ssRNA receptor, while pDCs express TLR7 (14). Consistent with this, CU-CPT9a blocked the production of pro-inflammatory cytokines in cDCs (Figure 5A). Our effort to block TLR7 signaling using commercially available receptor antagonists was unsuccessful since none of these inhibitors blocked TLR7 activation in pDCs stimulated with R848 or Imiquimod (data not shown). As an alternative strategy to demonstrate the involvement of TLR7 in SCV2-RNA sensing we performed TLR desensitization (21). pDCs were stimulated with SCV2-RNA or R848 or left untreated, washed, and then re-stimulated with R848. Figure 5B shows that, upon re-stimulation, only untreated cells could respond to R848 in terms of IFN-α and TNF-α production as a result of TLR7 desensitization by its ligand R848 as well as by SCV2-RNA. The limited yield following blood DC purification hampered the use of MyD88 siRNA. However, the involvement of endosomal TLRs as SCV2-RNA receptors was further supported by the blocking of cytokine release in both cDCs (Figure 5C) and pDCs (Figure 5D) by chloroquine (CQ), a drug known to block endosomal TLR triggering by interfering with endosomal acidification (27).

**Figure 4.**
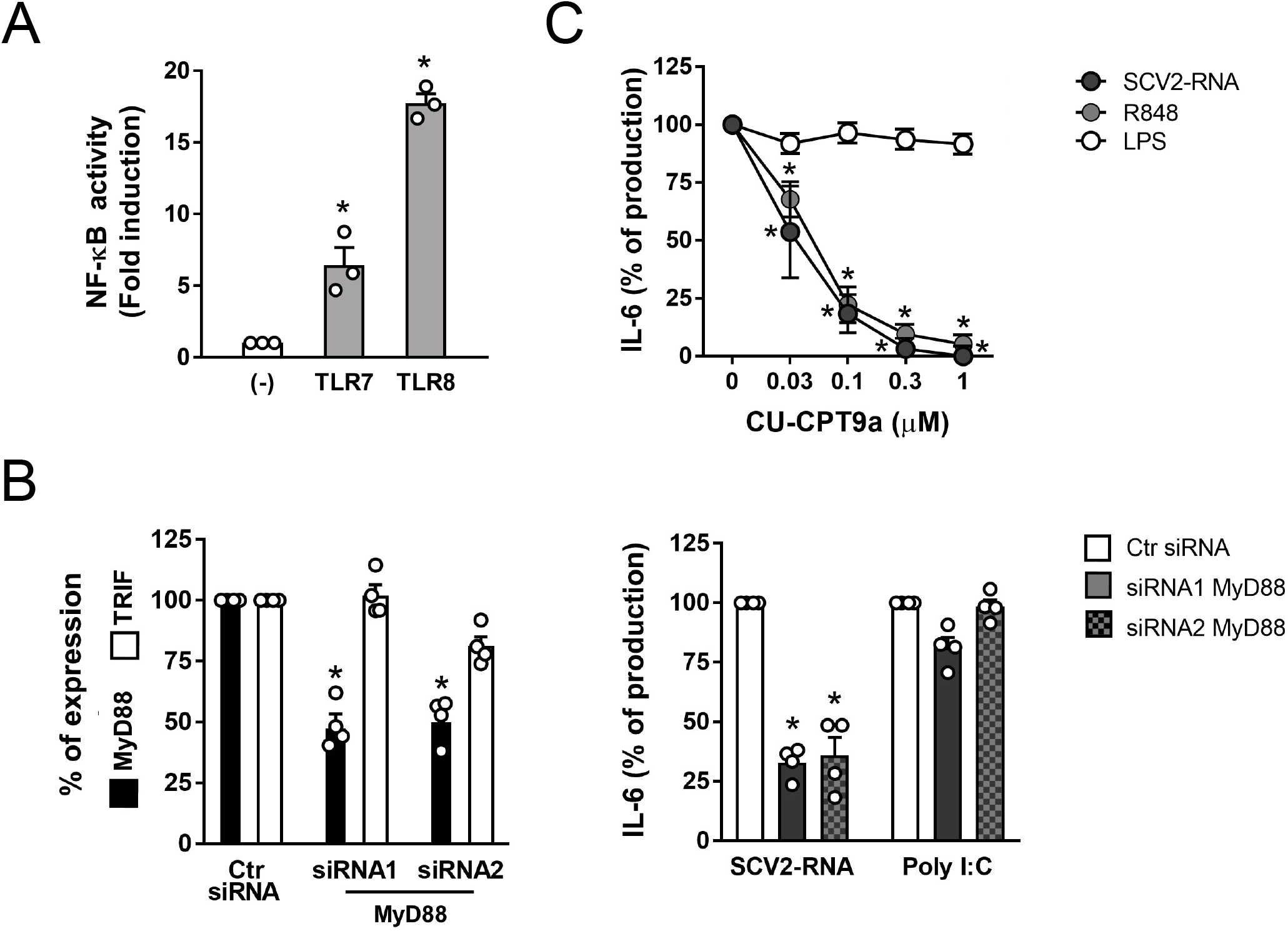
TLR7 and TLR8 are responsible for moDC activation by SAMPs. (A) Reporter HEK-293 cells stably transfected with human TLR7, TLR8 or luciferase alone (-) were stimulated with 10 μg/ml SCV2-RNA for 24 hours. NF-κB activation was evaluated in terms of luciferase activity. Data are expressed ad mean ± SEM (n=3); *P< 0.05 versus (-) by one-way ANOVA with Dunnett’s post-hoc test. (B, left panel) moDCs were transfected with MyD88-specific siRNAs or with control siRNA and the expression of MyD88 and TRIF was evaluated by Real-time PCR. Results are expressed as percentage of expression of MyD88 (black bars) and TRIF (white bars) in cell transfected with control siRNA (mean ± SEM n=4); *P< 0.05 versus respective “ctr siRNA” by one-way ANOVA with Dunnett’s post-hoc test. (B, right panel) moDCs transfected with MyD88-specific siRNAs or with control siRNA were stimulated with 5 μg/ml SCV2-RNA or 25 μg/ml Poly I:C for 24 hours. The production of IL-6 was evaluated by ELISA. Data are expressed as percentage of production for each individual stimulation (n=4); *P< 0.05 versus respective “ctr siRNA” by one-way ANOVA with Dunnett’s post-hoc test. (C) moDCs were pre-treated with increasing concentration of CU-CPT9a for 1 hour and then stimulated with SCV2-RNA (5 μg/ml) or R848 (1 μg/ml) or LPS (100 ng/ml) for 24 hours. IL-6 production was evaluated by ELISA. Data are expressed as percentage of production for each individual stimulation (n=3); *P< 0.05 versus respective “0” by one-way ANOVA with Dunnett’s post-hoc test.

**Figure 5.**
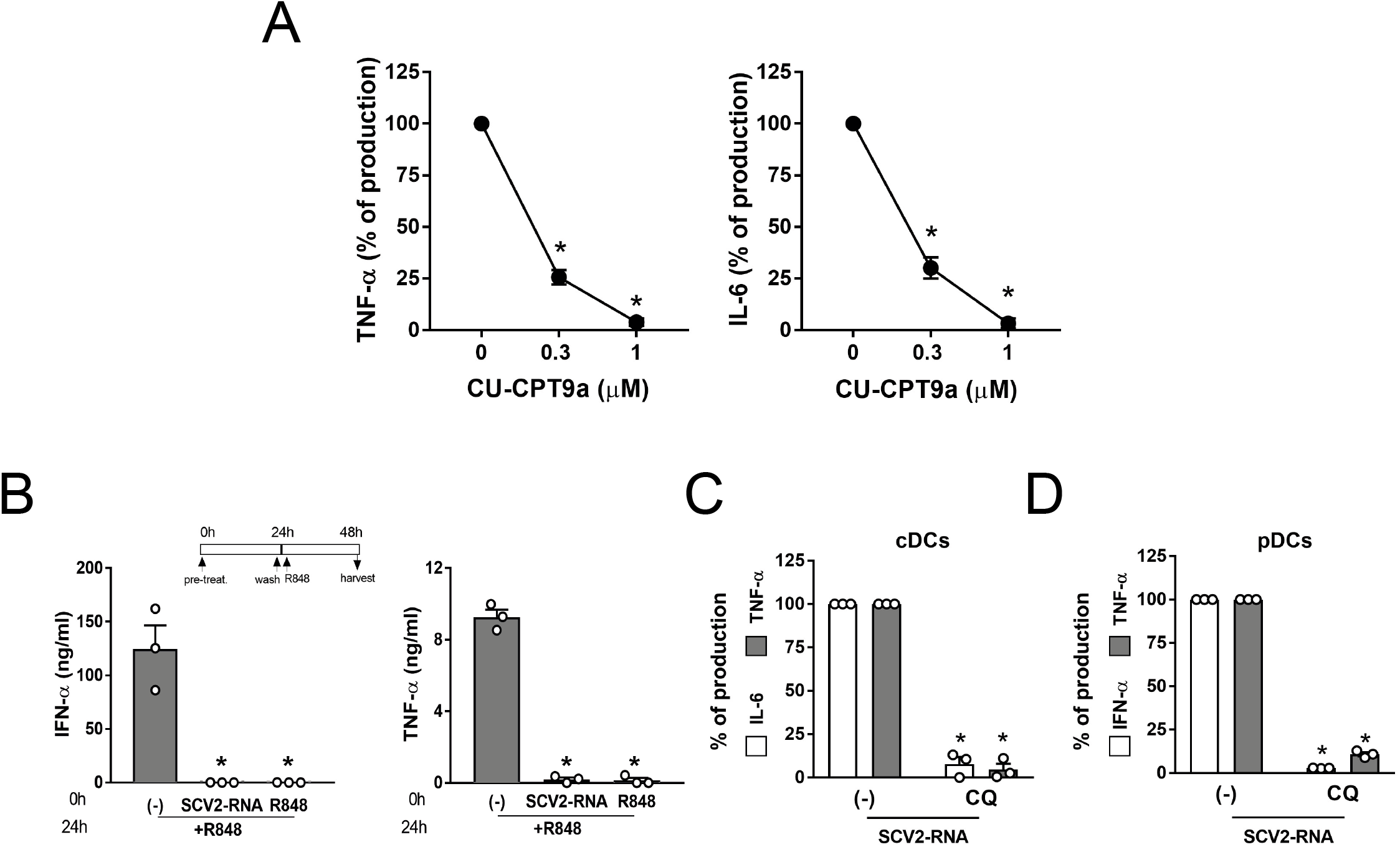
TLR7 and TLR8 are responsible for primary DC activation by SAMPs. (A) cDCs were pre-treated with increasing concentration of CU-CPT9a for 1 hour and then stimulated with SCV2-RNA (5 μg/ml) for 24 hours. Secreted TNF-α and IL-6 were quantified by ELISA. Data are expressed as percentage of production (n=3); *P< 0.05 versus “0” by one-way ANOVA with Dunnett’s post-hoc test. (B) pDCs were pre-treated (0h) with SCV2-RNA (5 μg/ml) or R848 (1 μg/ml) or left untreated for 24 hours, washed and restimulated with R848 for additional 24 hours. Secreted IFN-α and TNF-α were quantified by ELISA. Data are expressed as mean ± SEM (n=3); *P< 0.05 versus “(-)” by one-way ANOVA with Dunnett’s post-hoc test. (C) cDCs were pre-treated for 1 hour with Chloroquine (CQ, 10 μM) and then stimulated with SCV2-RNA (5 μg/ml) for 24 hours. Secreted IL-6 (white bars) and TNF-α (grey bars) were evaluated by ELISA. Data are expressed as percentage of production (n=3); *P< 0.05 versus respective “(-) SCV2-RNA” by paired Student’s *t* test. (D) pDCs were pretreated for 1 hour with CQ (10 μM) and then stimulated with SCV2-RNA (5 μg/ml) for 24 hours. Secreted IFN-α (white bars) and TNF-α (grey bars) were quantified by ELISA. Data are expressed as percentage of production (n=3); *P< 0.05 versus respective “(-) SCV2-RNA” by paired Student’s *t* test.

### ssRNA SAMPs induce DC activation and lung inflammation in vivo

To address the capacity of SAMPs to induce inflammation and immune activation *in vivo*, we first investigated if SAMPs can also trigger murine TLRs. TLR expression was analyzed in RAW264.7, a murine cell line, showing the expression of the ssRNA receptor TLR7 among other TLRs (Figure 6A). These TLR7-bearing cells responded to SAMP stimulation by producing TNF-α and this effect was blocked by CQ (Figure 6B) confirming that SCV2-RNA activate murine cells, presumably via TLR7. In addition, splenocytes from MyD88^−/−^ mice did not produce pro-inflammatory cytokines when stimulated with SCV2-RNA despite expressing similar levels of TLRs (Figure 6C and D). Based on these results, C57Bl6/J WT and MyD88^−/−^ mice were injected i.v. with SAMPs or vehicle and sacrificed 6 hours later. A significant increase of type I IFN was detected in the sera of WT SAMP-treated mice indicating systemic immune activation (Figure 6E). Consistent with this, SAMPs induced the upregulation of CD40 and CD86 on splenic pDCs (CD11c^int^MHC-II^+^B220^+^SiglecH^+^) (Figure 6F). Activation of splenic cDC1s (CD11c^+^MHC-II^+^CD8α^+^CD11b^−^) and cDC2s (CD11c^+^MHC-II^+^CD8α^−^CD11b^+^) was also detected (Figure 6G and H). Figure 7A shows that SAMP treatment induced the expression of pro-inflammatory cytokines TNF-α, IL-1β and IL-6 and of IFN-α and IFN-γ in the lung. In addition, a marked increase in the expression of chemokines active on myeloid and Th1 effector cells (i.e. CCL3, CCL4 and CXCL10) was also detected. Conversely, CCL20 and CCL22, two chemokines active in Th17 and Th2 T cell recruitment, were not increased (Figure 7B). We could also detect the accumulation of molecules involved in cytotoxic tissue damage such as GrB and TRAIL (Figure 7C) that, given the short kinetics of stimulation, may reflect the recruitment of NK cells to the lungs. The increase of CD45 and MHC-II mRNA levels (data not shown) further suggested immune cell infiltration, which was confirmed by histological analysis. Lung histology revealed a marked infiltration of inflammatory cells into peri-bronchial and peri-vascular connective tissue and alveolar septal thickening in SAMP-treated mice (Figure 7D). On the contrary, SAMP administration to MyD88^−/−^ mice did not induce any inflammatory response, including the increase of circulating levels of type I IFN, DC maturation and the generation of a lung infiltrate (Fig. 6 D-H and Figure 7). These data extend to the *in vivo* condition the observation that SAMPs use a TLR/MyD88-dependent pathway to trigger a type I IFN/pro-inflammatory activation program and highlight lung as a primary target organ.

**Figure 6.**
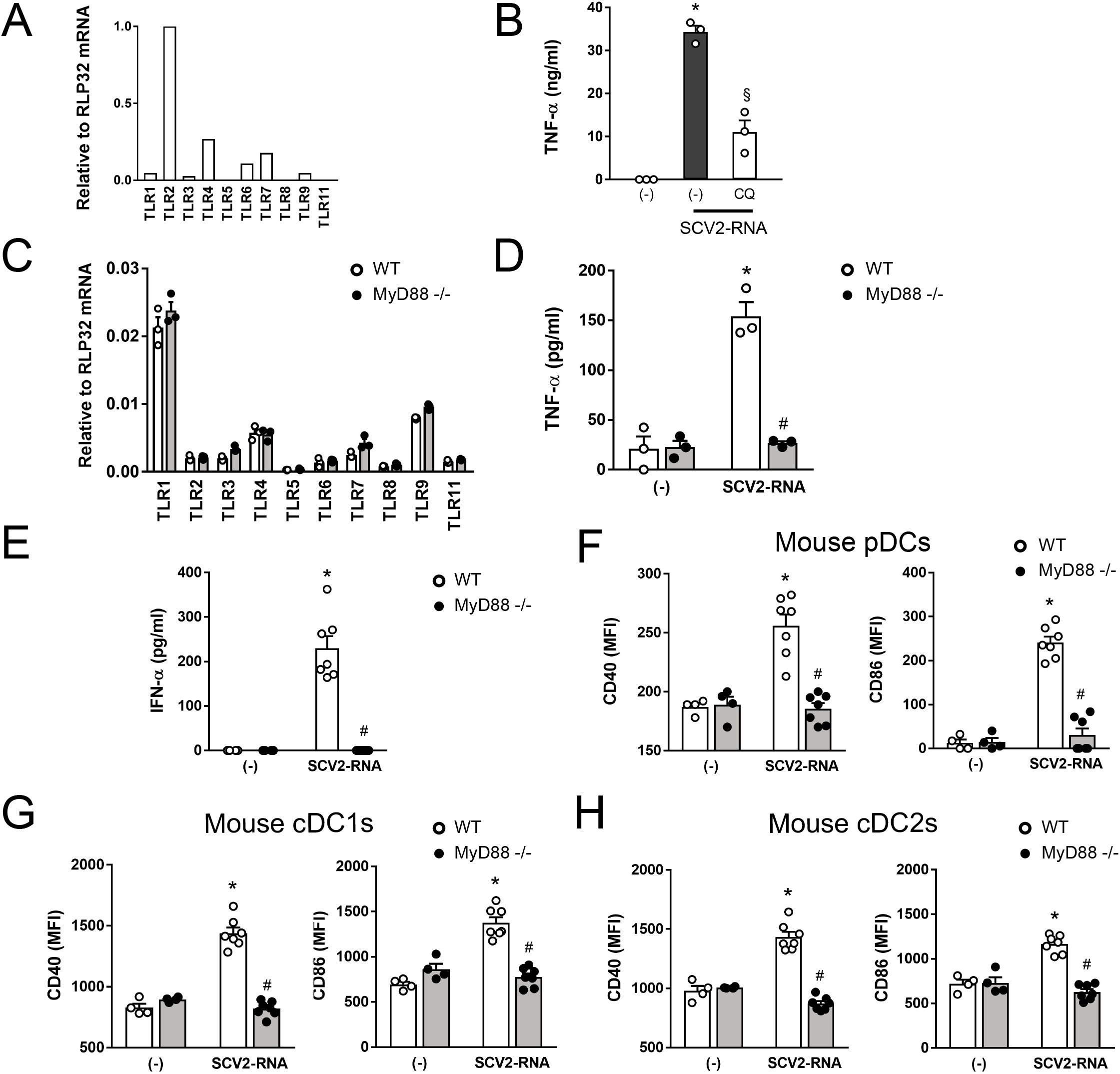
SAMPs activate murine cells *in vitro* and *in vivo*. (A) Expression of TLR mRNAs in RAW264.7 cells. Data are expressed as 2^−ΔCt^ relative to RPL32 of one representative experiment out of three. (B) RAW264.7 (1×10^6^/ml) were pre-treated for 1 hour with CQ (12.5 μM), then stimulated with 5 μg/ml SCV2-RNA or vehicle (-) for 24 hours. Secreted TNF-α was evaluated by ELISA. Data are expressed as mean ± SEM (n=3); *P< 0.05 versus (-); §P<0.05 versus “(-) SCV2-RNA” by paired Student’s *t* test. (C) Expression of TLR mRNAs in splenocytes from WT (white circle) or MyD88^−/−^ mice (black circle). Data are expressed as mean ± SEM (n=3) of 2^−ΔCt^ relative to RPL32 of one representative experiment out of three. (D) Splenocytes (3×10^6^/ml) from WT (white circle) or MyD88^−/−^ mice (black circle) were stimulated with 5 μg/ml SCV2-RNA or vehicle (-) for 24 hours. Secreted TNF-α was evaluated by ELISA. Data are expressed as mean ± SEM (n=3); *P< 0.05 versus (-) or #P < 0.05 versus “SCV2-RNA MyD88^−/−^” by paired Student’s *t* test. (E) Circulating IFN-α in WT (white circle) or MyD88^−/−^ mice (black circle) treated with SCV2-RNA or vehicle (-) for 6 hours. Data are expressed as mean ± SEM ((-) n=4, SCV2-RNA n=7); *P< 0.05 versus (-) or #P < 0.05 versus “SCV2-RNA MyD88^−/−^” by unpaired Student’s *t* test of one representative experiment out of three. (F-H) Activation of splenic pDCs (CD11c^int^MHC-II^+^B220^+^SiglecH^+^) (F), cDC1s (CD11c^+^MHC-II^+^CD8α^+^CD11b^−^) (G) or cDC2s (CD11c^+^MHC-II^+^CD8α^−^CD11b^+^) (H) from WT (white circle) or MyD88^−/−^ mice (black circle), treated with SCV2-RNA or vehicle (-) for 6 hours evaluated in terms of CD40 and CD86 expression. Data are expressed as mean ± SEM of the median fluorescence intensity (MFI) ((-) n=4, SCV2-RNA n=7); *P< 0.05 versus (-) or #P < 0.05 versus “SCV2-RNA MyD88^−/−^” by unpaired Student’s *t* test.

**Figure 7.**
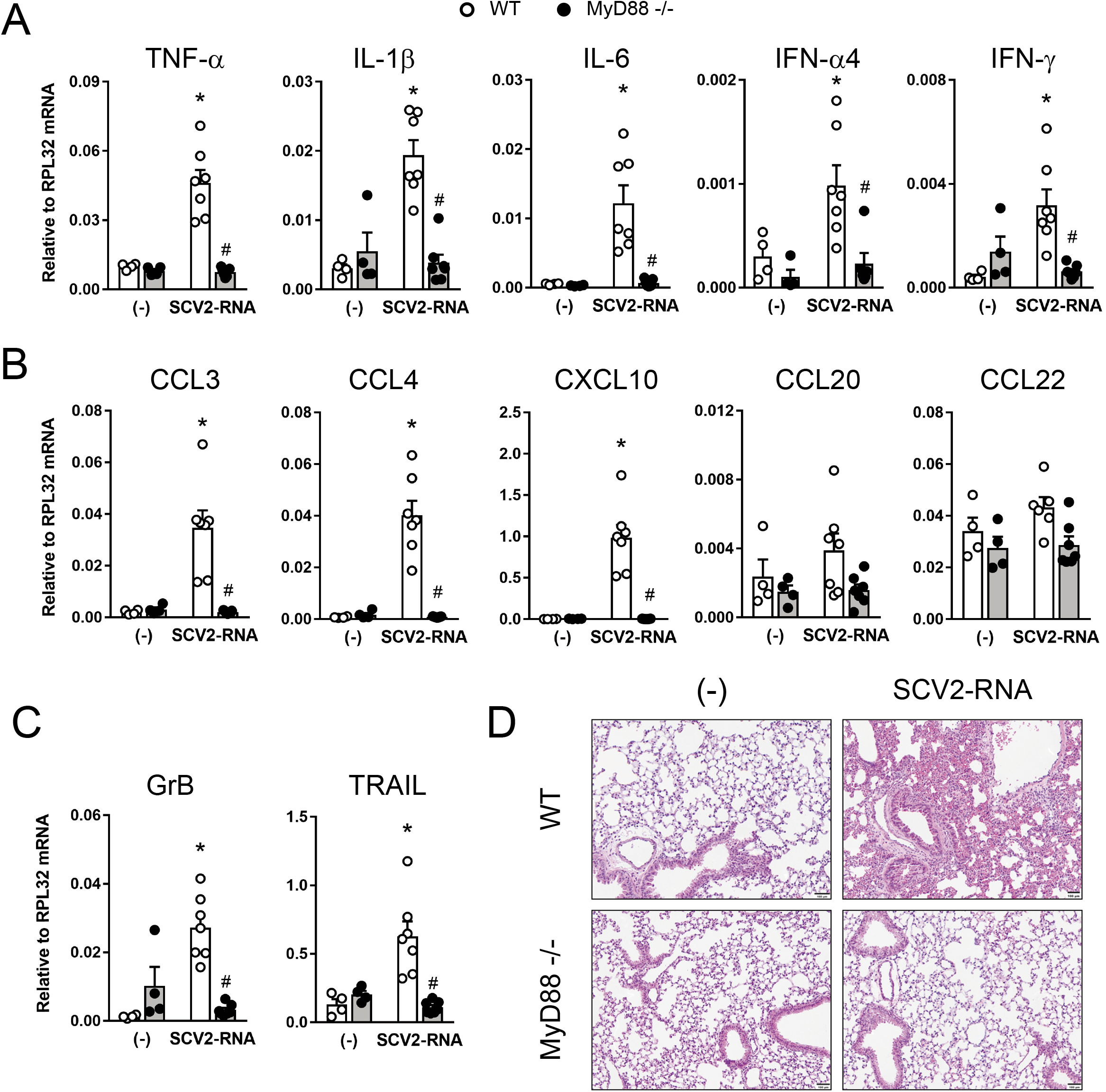
SAMPs induce inflammation *in vivo*. (A-C) Real-time PCR for cytokines, chemokines and effector proteins in lungs of WT (white circle) or MyD88^−/−^ (black circle) treated or not with SCV2-RNA for 6 hours. Data are expressed as mean ± SEM ((-) n=4, SCV2-RNA n=7) of 2^−ΔCt^ relative to housekeeping mRNA (RPL32); *P< 0.05 versus (-) or #P < 0.05 versus “SCV2-RNA MyD88^−/−^” by unpaired Student’s *t* test. (D) Histological evaluation of lungs from WT or MyD88^−/−^ mice treated or not with SCV2-RNA for 6 hours. One representative section is shown. Scale bars = 100 μm.

## Discussion

Here, we report that two short sequences within the ssRNA genome of SARS-CoV-2 activate the production of type I IFNs and the T cell-activating ability of human DCs by triggering endosomal TLR7 and TLR8. Of note, these sequences represent prototypical examples of the several hundreds of potential TLR ligands identified by SARS-CoV-2 genome scan. This finding is in line with previous work demonstrating a twenty-fold higher density of GU-rich fragments in the closely related SARS-CoV as compared to HIV-1 (28) and with a recent bioinformatic study showing that SARS-CoV-2 encodes a number of such fragments even larger than SARS-CoV (29). Thus, endosomal processing of SARS-CoV-2 nucleic acids may give rise to multiple fragments endowed with the property to trigger innate immune activation.

DCs play a crucial role as activators of both inflammation and adaptive immune responses (7, 8) and pDCs are the major producers of type I IFNs in response to viral infections (10–12). The protective role of type I IFN in life-threatening COVID-19 has been documented based on the clinical outcome of patients with inborn errors in type I IFN immunity or producing blocking auto-Abs against different types of type I IFNs (5, 6). Therefore, SAMPs may represent one of the essential signals in the activation of an IFN response and Th1-oriented adaptive immunity (30, 31). In this regards it is of note that SARS-CoV-2 infection affected the number of pDCs in vivo (17, 18) and primary virus isolates induced the activation of pDCs, *in vitro* (32). Here we extend this knowledge through the identification of viral sequences active on pDCs as well as cDCs and moDCs and show that the activation program induced by SAMPs is not restricted to type I IFNs, but encompassed the production of pro-inflammatory cytokines and the generation of Th1-oriented responses, supporting a possible role for these cells in the generation of the exuberant pro-inflammatory response observed in life-threatening COVID-19(33).

Very rare loss-of-function variants of TLR7 in two independent families was associated with severe COVID-19 in males (34). Thus, our report on the ability of SAMPs to activate the TLR7/8 and MyD88 pathways provides the missing link between this clinical evidence and our molecular knowledge on the cellular sensors for SARS-CoV-2 detection. Viral recognition by endosomal TLRs takes place before and independently of infection, as a consequence of pathogen endocytosis (13). Indeed, pDCs were reported to be resistant to infection, although they were activated by SARS-CoV-2 (32). This is an important process that gives innate immune cells the opportunity to activate early antiviral response. It remains to be elucidated if SARS-CoV-2 uptake for endosomal processing is a direct process or mediated by receptors, such as ACE2 or CD147 (35). ssRNA-sensing TLRs are expressed also by other innate immune cells, such as macrophages, as well as by peripheral tissues such as lung, bronchus, rectum, and cerebral cortex (36). Thus, other cells may support as well both protective and excessive antiviral responses (4). Since the magnitude of TLR activation differs in individuals, such as elderly people, differences in TLR activation may help explain differences in the quality of the antiviral immune response independently of SAMP potency (37).

By all means, other SAMPs and DAMPs as well as the simultaneous engagement of different PRRs are likely to contribute to COVID-19-associated protective response and cytokine storm, including cytosolic sensors, such as retinoid-inducible gene-1 (RIG1)-like receptors **(38)**, Interferon Induced proteins with tetratricopeptide repeats, or members of a large group of RNA-binding molecules with poorly defined ligand specificity **(38)**. A search for specific candidate ligands of cytosolic RNA-sensors was hampered because the scarce definition of their ligand consensus sequences. However, the finding that SARS-CoV-2 can evade innate immune restriction provided by intracellular RNA-sensors via methylation the 5’-end of its cellular mRNAs **(39)** further reinforces the role for TLRs as crucial sentinels and regulators of immune response to SARS-CoV-2 infection. SARS-CoV-2 is known to induce inflammasome assembly despite the exact mechanism still need to be characterized (40, 41). Since intracellular nucleic acid sensors are known to activate inflammasomes (42), and TLR activation is intimately connected with inflammasome functions (43), it is possible that SCV2-RNAs used in this study may also contribute to activate this pathway.

In conclusion, this work describes that SARS-CoV-2 is as a potential powerful source of immunostimulatory nucleic acid fragments and identifies the first SARS-CoV-2-specific PAMPs endowed with the ability to promote inflammation and immunity triggering TLR7 and TLR8. Based on previous works demonstrating a) the crucial protective role of type I IFNs against COVID-19 (5, 6); b) the crucial protective role of TLR7 against life-threatening SARS-CoV-2 infection (34) and c) pDC activation *in vitro* by SARS-CoV-2 (32), we believe that our findings fill a gap in the understanding of SARS-CoV-2 host-pathogen interaction.

## Methods

### Identification of potential TLR7/8-triggering ssRNA PAMPs

The reference SARS-CoV-2 genome (NC_045512, positive strand) was scanned for GU-rich ssRNA fragments with the SequenceSearcher tool in the Fuzzy mode (44). We defined “GU-enriched sequences” short strings with a maximal length of 20 bp, that were composed for more than 40% of the length by “GU” and/or “UG” pairs. The identified 491 GU-rich sequences were further selected based on the content of at least one “UGUGU” Interferon Induction Motif (IIM)(21) (see Suppl. Table 1). Within this list, the following were selected and synthesized by Integrated DNA Technologies (IDT) for subsequent studies: SCV2-RNA1 5’-UGCUGUUGUGUGUU*U-3’ (genome position: 15692-15706); SCV2-RNA2 5’-GUGUGUGUGUUCUGUUAUU*G-3’ (genome position: 20456-20475). These sequences were checked for uniqueness with BLAST in the database RefSeq Genome Database (refseq_genomes) within the RNA viruses (taxid: 2559587). Two additional sequences were synthesized, in which “U” was substituted with “A”, in order to impair TRL7/8 stimulation (SCV2-RNA1A and SCV2-RNA2A) (15, 20). * indicates a phosphorothioate linkage.

### Cell preparation and culture

Buffy coats from blood donations of anonymous healthy donors were obtained and preserved by the Centro Trasfusionale, Spedali Civili of Brescia according to the italian law concerning blood component preparation and analysis. Peripheral blood mononuclear cells (PBMC) were obtained by density gradient centrifugation and monocytes were subsequently purified by immunomagnetic separation using anti CD14-conjugated magnetic microbeads (Miltenyi Biotec) according to the manufacture’s protocol and as previously published (23). Briefly, monocytes were cultured for 6 days in tissue culture plates in complete medium (RPMI 1640 supplemented with 10%heat-inactivated, endotoxin free FBS, 2 mM L-Glutamine, penicillin and streptomycin (all from Gibco, Thermo Fisher Scientific) in the presence of 50 ng/ml GM-CSF and 20 ng/ml IL-4 (Miltenyi Biotec). Untouched peripheral blood cDC1 and cDC2 (cDCs) and pDCs were obtained from PBMC after negative immunomagnetic separation with the Myeloid Dendritic Cell Isolation kit (Miltenyi Biotec) and the Plasmacytoid Dendritic Cell Isolation kit II (Miltenyi Biotec), respectively. pDCs were cultured in completed RPMI medium with 20 ng/ml IL-3 (Miltenyi Biotec). RAW264.7 cells were purchased from American Type Culture Collection and cultured in DMEM complemented with 10% FBS.

### Cell stimulation with viral RNAs

Complexation of RNA with DOTAP Liposomal Transfection Reagent (Roche) was performed as previously described (21). Briefly, 5 μg RNA in 50 μl HBS buffer (20 mM HEPES, 150 mM NaCl, pH 7.4) was combined with 100 μl DOTAP solution (30 μl DOTAP plus 70 μl HBS buffer) and incubated for 15 minutes at RT. Where indicated, cells were pretreated for 1 hour with Chloroquine or CU-CPT9a or stimulated with TLR agonists (all from Invivogen).

### MyD88 silencing in moDCs

Differentiating monocytes at day 2 of culture were transfected with two different MyD88 Silencer Select Validated siRNA or with a control siRNA (all at 50 nM final concentration; Ambion, Thermo Fisher Scientific) using Opti-MEM I reduced serum medium and Lipofectamine RNAiMAX transfection reagent (Thermo Fisher Scientific) as previously described (45). Transfected cells were incubated for 72 hours and then stimulated for 24 hours with TLR agonists as indicated. The effects of mRNA silencing by siRNA was investigated by real-time PCR using specific QuantiTect primer Assay (Qiagen).

### Cytokine detection

TNF-α, IL-6, IL-12p70, CXCL8, CXCL9, CCL3 and mouse TNF-α were measured by ELISA assay (R&D Systems). Human IFN-α was detected using specific Module Set ELISA kit (eBioscience). Mouse IFN-α was measured by a bioluminescence kit (InvivoGen). All assays were performed on cell free supernatants according to the manufacturer’s protocol.

### Flow cytometry

Human and mouse DCs were stained with the following antibodies from Miltenyi Biotec or as specified: Vioblue-conjugated anti-human CD86 (clone FM95, Miltenyi Biotec), PE-conjugated anti-human CD83 (clone REA714), FITC-conjugated anti-human BDCA2 (clone AC144), APC-conjugated anti-human CCR7 (clone REA546), VioGreen-conjugated anti-mouse CD45 (clone REA737), VioBlue or FITC-conjugated anti-mouse MHCII (clone REA564), PerCP-Vio 700-conjugated anti-mouse CD11c (clone REA754), PE-conjugated anti-mouse SiglecH (clone 551.3D3), PE-Vio 615-conjugated anti-mouse CD11b (clone REA592), VioBlue-conjugated anti-mouse CD8a (cloneREA601), PE-Vio 770-conjugated anti-mouse B220 (clone RA3-6B2), PE-conjugated anti-mouse CD40 (clone REA965), FITC-conjugated anti-mouse CD40 (clone HM40-3, Biolegend) and APC-CY7-conjugated anti-mouse CD86 (clone GL-1, Biolegend). Samples were read on a MACSQuant Analyzer (Miltenyi Biotec) and analysed with FlowJo (Tree Star Inc.). For intracellular detection of Granzyme B, cells were fixed and permeabilized using the Inside Stain kit (Miltenyi Biotec) and stained with APC-conjugated anti-Granzyme B (clone REA226, Miltenyi Biotec). Cell viability was assessed by LIVE/DEAD staining according to the manufacturer’s instruction (Molecular Probes, Thermo Fisher Scientific).

### NF-κB luciferase reporter assay

TLR-specific activation assays were performed using human HEK293 cells expressing luciferase under control of the NF-κB promoter and stably transfected with human TLR7 and TLR8 as previously described (21). Briefly, 25000 cells were seeded in complete DMEM without antibiotics in 96-well plates for 24 hours and then stimulated with 10 μg/ml SCV2-RNA for additional 24 hours. After stimulation, cells were lysed using ONE-Glo EX Luciferase Assay System (Promega) according to the manufacturer’s recommendations and assayed for luciferase activity using the EnSightMultimode Plate Reader (PerkinElmer). HEK293-transfected cells were maintained in DMEM supplemented with 10% FBS and specific antibiotics were added.

### T cell proliferation assay

Experiments using T cells were conducted according to the “Minimal Information about T Cell Assays” (MIATA) guidelines. Allogenic naïve CD4+ T cells and CD8+ T cells were isolated from buffycoats using the naïve CD4+ T cell Isolation kit II (Miltenyi Biotec) and CD8+ T cell Isolation kit (Miltenyi Biotec), respectively. Purified T cells were counted by flow cytometry and labeled with CellTrace-CFSE (Molecular Probes, Thermo Fisher Scientific) at a final concentration of 5 μM. Subsequently, T cells (1×105 cells/well) were cocultured with graded numbers of allogeneic moDCs in 96-well round-bottom culture plates in complete RPMI medium. After 6 days, alloreactive T cell proliferation was assessed by measuring the loss of the dye CellTrace-CFSE upon cell division using flow cytometry. Positive controls of T cell proliferations were routinely performed using IL-2 plus PHA. Response definition criteria were defined post-hoc. Dead cells were excluded by LIVE/DEAD staining according to the manufacturer’s instruction. These experiments were performed using general research investigative assays. Raw data can be provided per request.

### Analysis of T cell cytokine production

After 6 days of coculture, helper T cells were restimulated with 200 nM PMA (Sigma-Aldrich) plus 1 μg/ml of ionomycin (Sigma) for 5 hours. Brefeldin A (5 μg/ml, Sigma) was added during the last 2 hours. For intracellular cytokine production, cells were fixed and permeabilized with Inside Stain kit (Miltenyi Biotec) and stained with FITC-conjugated anti-IFN-γ (clone 45-15, Miltenyi Biotec) and PE-conjugated anti-IL-4 (clone 7A3-3, Miltenyi Biotec) following the manufacturer’s recommendations. For CD8^+^ T cells, after 6 days of coculture, IFN-γ production was assessed in the culture supernatants by ELISA (R&D system). Response definition criteria were defined post-hoc. These experiments were performed using general research investigative assays. Raw data can be provided per request.

### *In vivo* experiments

Procedures involving animal handling and care conformed to protocols approved by the University of Verona in compliance with national (D.L. N.116, G.U., suppl. 40, 18-2-1992 and N. 26, G.U. March 4, 2014) and international law and policies (EEC Council Directive 2010/63/EU, OJ L 276/33, 22-09-2010; National Institutes of Health Guide for the Care and Use of Laboratory Animals, U.S. National Research Council, 2011). The study was approved by the Italian Ministry of Health (approval number 339/2015-PR). Sex and age matched C57Bl6/J mice were obtained by Charles River Laboratories and housed in the specific pathogen-free animal facility of the Department of Medicine, University of Verona. MyD88^−/−^ mice were kindly provided by S. Akira (Osaka University, Osaka, Japan). Mice were anesthetized with isoflurane and injected i.v. in the retro-orbital vein with 300 μl DOTAP/SCV2-RNA mixture (20 μg/mouse) or with DOTAP alone. After 6 hours, mice were sacrificed and lungs, spleen and blood were harvested. Briefly, lungs were collected upon intracardiac perfusion with cold PBS. Left lung lobes were formalin fixed for 24 hours, dehydrated, and paraffin embedded for histological analysis. Right lungs were immediately frozen at −80°C and used for real-time PCR. Spleens were mechanically and enzymatically treated to obtain a single-cell suspension for cytofluorimetric and real-time PCR analysis. All mouse experiments were carried out in accordance with guidelines prescribed by the Ethics Committee for the use of laboratory animals for research purposes at the University of Verona and by the Italian Ministry of Health. All efforts were made to minimize the number of animals used and their suffering.

### Lung histological analysis

Histology was performed on three longitudinal serial sections (150 μm apart, 4 μm in thickness) from each left lung, stained with hematoxylin and eosin (H&E), and scanned by VS120 Dot-Slide BX61 virtual slide microscope (Olympus Optical) as previously described (46).

### Real-time PCR

RNA was extracted using TRIzol reagent, treated with DNAse according to the manufacturer’s instructions and reverse transcription performed using random hexamers and MMLV RT (all from Thermo Fisher Scientific). The SsoAdvanced Universal SYBR Green Supermix (Bio-Rad Laboratories) was used according to the manufacturer’s instructions. Reactions were run in triplicate on a StepOne Plus Real-Time PCR System (Applied Biosystems) and analyzed by the StepOne Plus Software (Version 2.3, Applied Biosystems). Sequences of gene-specific primers are available upon request. Gene expression was normalized based on RPL32 mRNA content.

### Statistical analysis

Statistical significance among the experimental groups was determined using paired or unpaired Student’s *t* test or one-way ANOVA with Dunnet’s post-hoc test (GraphPad Prism 7, GraphPad Software) as indicated in each figure legend. P< 0.05 was considered significant.

## Author Contributions

Conceptualization, D.B., M.A.C., A.M., S.S., V.S., A.D.P.; methodology, V.S., A.D.P., P.S., D.B., C.G.; software, M.L. and D.B.; validation and formal analysis, V.S., H.O.N. M.P., I.B.; investigation, V.S., H.O.N., F.S., T.S., M.L., P.S., L.T., I.B., M.P, N.T., C.G.; data curation, V.S., H.O.N., F.S., M.L., I.B.; writing—original draft preparation, D.B., V.S., H.O.N.; writing—review and editing, A.M., S.S., M.A.C., P.S., A.D.P, D.B.; visualization, H.O.N. and I.B.; supervision, D.B., M.A.C., S.S.; funding acquisition, S.S., M.A.C, D.B., P.S. All authors have read and agreed to the published version of the manuscript.

## Acknowledgements

This research was funded by the Italian Ministry of Health (Bando Ricerca COVID-2020-12371735 to S.S.), Italian Ministry of the University and Research (MUR-PRIN 20178ALPCM_005 to D.B. and MUR-PRIN 20177J4E75_004 to M.A.C.), Associazione Italiana per la Ricerca sul Cancro (AIRC IG-20339 to M.A.C. and IG-20776 to S.S.), University of Verona (RBVR17NCNC to P.S.) and University of Brescia (Fondi Locali 2019 and 2020 to D.B.). A.M. acknowledges support by the Dolce & Gabbana fashion company.

