## Supplemental Table 1 for "SARS-CoV-2-associated ssRNAs activate inflammation and immunity via TLR7/8"

| Sequence number | Sequence | Start | Stop |
|---|---|---|---|
| 1 | TTTAAAATC**TGTGT**GGCTG | 76 | 94 |
| 2 | TAAAATC**TGTGT**GGCTGTC | 78 | 96 |
| 3 | C**TGTGT**GGCTGTCACTCGG | 84 | 102 |
| 4 | TTGCCTTTGGAGGC**TGTGT** | 1476 | 1494 |
| 5 | TGCCTTTGGAGGC**TGTGT**G | 1477 | 1495 |
| 6 | GCCTTTGGAGGC**TGTGTGT** | 1478 | 1496 |
| 7 | CCTTTGGAGGC**TGTGTGT**T | 1479 | 1497 |
| 8 | CTTTGGAGGC**TGTGTGT**TC | 1480 | 1498 |
| 9 | TTTGGAGGC**TGTGTGT**TCT | 1481 | 1499 |
| 10 | TGGAGGC**TGTGTGT**TCTCT | 1483 | 1501 |
| 11 | GAGGC**TGTGTGT**TCTCTTA | 1485 | 1503 |
| 12 | GGC**TGTGTGT**TCTCTTATG | 1487 | 1505 |
| 13 | GC**TGTGTGT**TCTCTTATGT | 1488 | 1506 |
| 14 | C**TGTGTGT**TCTCTTATGTT | 1489 | 1507 |
| 15 | **TGTGTGT**TCTCTTATGTTG | 1490 | 1508 |
| 16 | G**TGTGT**TCTCTTATGTTGG | 1491 | 1509 |
| 17 | **TGTGT**TCTCTTATGTTGGT | 1492 | 1510 |
| 18 | TAAATTTTTGGCTT**TGTGT** | 2314 | 2332 |
| 19 | AAATTTTTGGCTT**TGTGT**G | 2315 | 2333 |
| 20 | AATTTTTGGCTT**TGTGT**GC | 2316 | 2334 |
| 21 | ATTTTTGGCTT**TGTGT**GCT | 2317 | 2335 |
| 22 | TTTTTGGCTT**TGTGT**GCTG | 2318 | 2336 |
| 23 | TTTTGGCTT**TGTGT**GCTGA | 2319 | 2337 |
| 24 | TTGGCTT**TGTGT**GCTGACT | 2321 | 2339 |
| 25 | GGCTT**TGTGT**GCTGACTCT | 2323 | 2341 |
| 26 | TT**TGTGT**GCTGACTCTATC | 2326 | 2344 |
| 27 | **TGTGT**GCTGACTCTATCAT | 2328 | 2346 |
| 28 | ATTGTACAGAAAG**TGTGT**T | 2410 | 2428 |
| 29 | AATGAGTTCGCC**TGTGT**TG | 2870 | 2888 |
| 30 | ATGAGTTCGCC**TGTGT**TGT | 2871 | 2889 |
| 31 | TGAGTTCGCC**TGTGT**TGTG | 2872 | 2890 |
| 32 | GAGTTCGCC**TGTGT**TGTGG | 2873 | 2891 |
| 33 | GTTCGCC**TGTGT**TGTGGCA | 2875 | 2893 |
| 34 | CC**TGTGT**TGTGGCAGATGC | 2880 | 2898 |
| 35 | **TGTGT**TGTGGCAGATGCTG | 2882 | 2900 |
| 36 | AAGTGGGTGGTAGT**TGTGT** | 3558 | 3576 |
| 37 | AGTGGGTGGTAGT**TGTGT**T | 3559 | 3577 |
| 38 | GTGGGTGGTAGT**TGTGT**TT | 3560 | 3578 |
| 39 | TGGGTGGTAGT**TGTGT**TTT | 3561 | 3579 |
| 40 | GGGTGGTAGT**TGTGT**TTTA | 3562 | 3580 |
| 41 | GGTGGTAGT**TGTGT**TTTAA | 3563 | 3581 |
| 42 | GTGGTAGT**TGTGT**TTTAAG | 3564 | 3582 |
| 43 | TGGTAGT**TGTGT**TTTAAGC | 3565 | 3583 |
| 44 | GTAGT**TGTGT**TTTAAGCGG | 3567 | 3585 |
| 45 | GT**TGTGT**TTTAAGCGGACA | 3570 | 3588 |
| 46 | ATTCTTTAAGAGTT**TGTGT** | 3744 | 3762 |
| 47 | TTCTTTAAGAGTT**TGTGT**A | 3745 | 3763 |
| 48 | CTTTAAGAGTT**TGTGT**AGA | 3747 | 3765 |
| 49 | TTAAGAGTT**TGTGT**AGATA | 3749 | 3767 |
| 50 | AAGAGTT**TGTGT**AGATACT | 3751 | 3769 |
| 51 | GAGTT**TGTGT**AGATACTGT | 3753 | 3771 |

| | | | |
|---|---|---|---|
| 52 | AGTT**TGTGT**AGATACTGTT | 3754 | 3772 |
| 53 | GTT**TGTGT**AGATACTGTTC | 3755 | 3773 |
| 54 | TT**TGTGT**AGATACTGTTCG | 3756 | 3774 |
| 55 | T**TGTGT**AGATACTGTTCGC | 3757 | 3775 |
| 56 | ATTAATGCCTGTC**TGTGT**G | 4426 | 4444 |
| 57 | TAATGCCTGTC**TGTGT**GGA | 4428 | 4446 |
| 58 | ATGCCTGTC**TGTGT**GGAAA | 4430 | 4448 |
| 59 | GCCTGTC**TGTGT**GGAAACT | 4432 | 4450 |
| 60 | AGTCTTGAACGTGG**TGTGT** | 5503 | 5521 |
| 61 | TCTTGAACGTGG**TGTGT**AA | 5505 | 5523 |
| 62 | CGTGG**TGTGT**AAAACTTGT | 5512 | 5530 |
| 63 | GTGGTGTGTAAAACTTGTG | 5513 | 5531 |
| 64 | TGG**TGTGT**AAAACTTGTGG | 5514 | 5532 |
| 65 | GG**TGTGT**AAAACTTGTGGA | 5515 | 5533 |
| 66 | G**TGTGT**AAAACTTGTGGAC | 5516 | 5534 |
| 67 | TTTATTGCTACAAT**TGTGT** | 6787 | 6805 |
| 68 | TTATTGCTACAAT**TGTGT**A | 6788 | 6806 |
| 69 | ATCAACTTGTATGA**TGTGT** | 7483 | 7501 |
| 70 | CAACTTGTATGA**TGTGT**TA | 7485 | 7503 |
| 71 | AATTGGAAT**TGTGT**TAATT | 7610 | 7628 |
| 72 | TTGGAAT**TGTGT**TAATTGT | 7612 | 7630 |
| 73 | GGAAT**TGTGT**TAATTGTGA | 7614 | 7632 |
| 74 | GAAT**TGTGT**TAATTGTGAT | 7615 | 7633 |
| 75 | AAT**TGTGT**TAATTGTGATA | 7616 | 7634 |
| 76 | AT**TGTGT**TAATTGTGATAC | 7617 | 7635 |
| 77 | **TGTGT**TAATTGTGATACAT | 7619 | 7637 |
| 78 | TTAAAGTTACACT**TGTGT**T | 8586 | 8604 |
| 79 | TTACACT**TGTGT**TCCTTTT | 8592 | 8610 |
| 80 | ACACT**TGTGT**TCCTTTTTG | 8594 | 8612 |
| 81 | CACT**TGTGT**TCCTTTTTGT | 8595 | 8613 |
| 82 | ACT**TGTGT**TCCTTTTTGTT | 8596 | 8614 |
| 83 | CT**TGTGT**TCCTTTTTGTTG | 8597 | 8615 |
| 84 | T**TGTGT**TCCTTTTTGTTGC | 8598 | 8616 |
| 85 | **TGTGT**TCCTTTTTGTTGCT | 8599 | 8617 |
| 86 | ACATCAGCT**TGTGT**TTTGG | 8993 | 9011 |
| 87 | ATCAGCT**TGTGT**TTTGGCT | 8995 | 9013 |
| 88 | CAGCT**TGTGT**TTTGGCTGC | 8997 | 9015 |
| 89 | AGCT**TGTGT**TTTGGCTGCT | 8998 | 9016 |
| 90 | GCT**TGTGT**TTTGGCTGCTG | 8999 | 9017 |
| 91 | CT**TGTGT**TTTGGCTGCTGA | 9000 | 9018 |
| 92 | T**TGTGT**TTTGGCTGCTGAA | 9001 | 9019 |
| 93 | **TGTGT**TTTGGCTGCTGAAT | 9002 | 9020 |
| 94 | AGAAGCTGGTGTT**TGTGT**A | 9241 | 9259 |
| 95 | AAGCTGGTGTT**TGTGT**ATC | 9243 | 9261 |
| 96 | GCTGGTGTT**TGTGT**ATCTA | 9245 | 9263 |
| 97 | TGGTGTT**TGTGT**ATCTACT | 9247 | 9265 |
| 98 | GTGTT**TGTGT**ATCTACTAG | 9249 | 9267 |
| 99 | GTT**TGTGT**ATCTACTAGTG | 9251 | 9269 |
| 100 | TT**TGTGT**ATCTACTAGTGG | 9252 | 9270 |
| 101 | T**TGTGT**ATCTACTAGTGGT | 9253 | 9271 |
| 102 | **TGTGT**ATCTACTAGTGGTA | 9254 | 9272 |
| 103 | TAGATTATGAC**TGTGT**CTC | 10509 | 10527 |
| 104 | GATTATGAC**TGTGT**CTCTT | 10511 | 10529 |

| | | | |
|---|---|---|---|
| 105 | TTATGAC**TGTGT**CTCTTTT | 10513 | 10531 |
| 106 | ATGAC**TGTGT**CTCTTTTTG | 10515 | 10533 |
| 107 | TGAC**TGTGT**CTCTTTTTGT | 10516 | 10534 |
| 108 | GAC**TGTGT**CTCTTTTTGTT | 10517 | 10535 |
| 109 | AC**TGTGT**CTCTTTTTGTTA | 10518 | 10536 |
| 110 | C**TGTGT**CTCTTTTTGTTAC | 10519 | 10537 |
| 111 | **TGTGT**CTCTTTTTGTTACA | 10520 | 10538 |
| 112 | TGCCGTTTTAGATA**TGTGT** | 10831 | 10849 |
| 113 | GCCGTTTTAGATA**TGTGT**G | 10832 | 10850 |
| 114 | CCGTTTTAGATA**TGTGT**GC | 10833 | 10851 |
| 115 | CGTTTTAGATA**TGTGT**GCT | 10834 | 10852 |
| 116 | GTTTTAGATA**TGTGT**GCTT | 10835 | 10853 |
| 117 | TTTTAGATA**TGTGT**GCTTC | 10836 | 10854 |
| 118 | TTAGATA**TGTGT**GCTTCAT | 10838 | 10856 |
| 119 | AGATA**TGTGT**GCTTCATTA | 10840 | 10858 |
| 120 | AC**TGTGT**TATGTATGCATC | 11307 | 11325 |
| 121 | **TGTGT**TATGTATGCATCAG | 11309 | 11327 |
| 122 | AGAAC**TGTGT**ATGATGATG | 11357 | 11375 |
| 123 | AAC**TGTGT**ATGATGATGGT | 11359 | 11377 |
| 124 | AC**TGTGT**ATGATGATGGTG | 11360 | 11378 |
| 125 | C**TGTGT**ATGATGATGGTGC | 11361 | 11379 |
| 126 | **TGTGT**ATGATGATGGTGCT | 11362 | 11380 |
| 127 | AGGTATTGTTTTTA**TGTGT** | 11533 | 11551 |
| 128 | GGTATTGTTTTTA**TGTGT**G | 11534 | 11552 |
| 129 | GTATTGTTTTTA**TGTGTGT** | 11535 | 11553 |
| 130 | TATTGTTTTTA**TGTGTGT**T | 11536 | 11554 |
| 131 | ATTGTTTTTA**TGTGTGT**TG | 11537 | 11555 |
| 132 | TTGTTTTTA**TGTGTGT**TGA | 11538 | 11556 |
| 133 | TGTTTTTA**TGTGTGT**TGAG | 11539 | 11557 |
| 134 | GTTTTTA**TGTGTGT**TGAGT | 11540 | 11558 |
| 135 | TTTTA**TGTGTGT**TGAGTAT | 11542 | 11560 |
| 136 | TTTA**TGTGTGT**TGAGTATT | 11543 | 11561 |
| 137 | TTA**TGTGTGT**TGAGTATTG | 11544 | 11562 |
| 138 | TA**TGTGTGT**TGAGTATTGC | 11545 | 11563 |
| 139 | **TGTGTGT**TGAGTATTGCCC | 11547 | 11565 |
| 140 | **TGTGT**TGAGTATTGCCCTA | 11549 | 11567 |
| 141 | ATTGTGGGCTCAA**TGTGT**C | 11923 | 11941 |
| 142 | TTGTGGGCTCAA**TGTGT**CC | 11924 | 11942 |
| 143 | TGTGGGCTCAA**TGTGT**CCA | 11925 | 11943 |
| 144 | TGGGCTCAA**TGTGT**CCAGT | 11927 | 11945 |
| 145 | TGCAAGAGATGGT**TGTGT**T | 12418 | 12436 |
| 146 | AGATGGT**TGTGT**TCCCTTG | 12424 | 12442 |
| 147 | ATGGT**TGTGT**TCCCTTGAA | 12426 | 12444 |
| 148 | GT**TGTGT**TCCCTTGAACAT | 12429 | 12447 |
| 149 | CTAAT**TGTGT**TAAGATGTT | 13140 | 13158 |
| 150 | TAAT**TGTGT**TAAGATGTTG | 13141 | 13159 |
| 151 | AAT**TGTGT**TAAGATGTTGT | 13142 | 13160 |
| 152 | AT**TGTGT**TAAGATGTTGTG | 13143 | 13161 |
| 153 | T**TGTGT**TAAGATGT**TGTGT** | 13144 | 13162 |
| 154 | **TGTGT**TAAGATGT**TGTGT**A | 13145 | 13163 |
| 155 | GTGTTAAGATGT**TGTGT**AC | 13146 | 13163 |
| 156 | AAT**TGTGT**TAACTGTTTGG | 14329 | 14347 |
| 157 | AT**TGTGT**TAACTGTTTGGA | 14330 | 14348 |

| | | | |
|---|---|---|---|
| 158 | T**TGTGT**TAACTGTTTGGAT | 14331 | 14349 |
| 159 | **TGTGT**TAACTGTTTGGATG | 14332 | 14350 |
| 160 | TTTAAGGAATTACT**TGTGT** | 14542 | 14560 |
| 161 | GGAATTACT**TGTGT**ATGCT | 14547 | 14565 |
| 162 | AATTACT**TGTGT**ATGCTGC | 14549 | 14567 |
| 163 | TTACT**TGTGT**ATGCTGCTG | 14551 | 14569 |
| 164 | ACT**TGTGT**ATGCTGCTGAC | 14553 | 14571 |
| 165 | TTCTATGACTTTGC**TGTGT** | 14695 | 14713 |
| 166 | CTATGACTTTGC**TGTGT**CT | 14697 | 14715 |
| 167 | TGACTTTGC**TGTGT**CTAAG | 14700 | 14718 |
| 168 | ACTTTGC**TGTGT**CTAAGGG | 14702 | 14720 |
| 169 | TTTGC**TGTGT**CTAAGGGTT | 14704 | 14722 |
| 170 | TTGC**TGTGT**CTAAGGGTTT | 14705 | 14723 |
| 171 | TGC**TGTGT**CTAAGGGTTTC | 14706 | 14724 |
| 172 | GC**TGTGT**CTAAGGGTTTCT | 14707 | 14725 |
| 173 | C**TGTGT**CTAAGGGTTTCTT | 14708 | 14726 |
| 174 | **TGTGT**CTAAGGGTTTCTTT | 14709 | 14727 |
| 175 | GTGAAATGGTCA**TGTGT**GG | 15431 | 15449 |
| 176 | AAATGGTCA**TGTGT**GGCGG | 15434 | 15452 |
| 177 | ATGGTCA**TGTGT**GGCGGTT | 15436 | 15454 |
| 178 | GGTCA**TGTGT**GGCGGTTCA | 15438 | 15456 |
| 179 | TCTGACGATGCTGT**TGTGT** | 15715 | 15733 |
| 180 | CTGACGATGCTGT**TGTGT**G | 15716 | 15734 |
| 181 | TGACGATGCTGT**TGTGTGT** | 15717 | 15735 |
| 182 | GACGATGCTGT**TGTGTGT**T | 15718 | 15736 |
| 183 | ACGATGCTGT**TGTGTGT**TT | 15719 | 15737 |
| 184 | CGATGCTGT**TGTGTGT**TTC | 15720 | 15738 |
| 185 | ATGCTGT**TGTGTGT**TTCAA | 15722 | 15740 |
| 186 | GCTGT**TGTGTGT**TTCAATA | 15724 | 15742 |
| 187 | TGT**TGTGTGT**TTCAATAGC | 15726 | 15744 |
| 188 | G**TGTGT**TTCAATAGCACTT | 15730 | 15748 |
| 189 | CAGGGTGATGATTA**TGTGT** | 15904 | 15922 |
| 190 | AGGGTGATGATTA**TGTGT**A | 15905 | 15923 |
| 191 | GGGTGATGATTA**TGTGT**AC | 15906 | 15924 |
| 192 | GGTGATGATTA**TGTGT**ACC | 15907 | 15925 |
| 193 | TGATGATTA**TGTGT**ACCTT | 15909 | 15927 |
| 194 | GATTA**TGTGT**ACCTTCCTT | 15913 | 15931 |
| 195 | AGGCTGTTGGGGCT**TGTGT** | 16235 | 16253 |
| 196 | GGCTGTTGGGGCT**TGTGT**T | 16236 | 16254 |
| 197 | GCTGTTGGGGCT**TGTGT**TC | 16237 | 16255 |
| 198 | CTGTTGGGGCT**TGTGT**TCT | 16238 | 16256 |
| 199 | TGTTGGGGCT**TGTGT**TCTT | 16239 | 16257 |
| 200 | GTTGGGGCT**TGTGT**TCTTT | 16240 | 16258 |
| 201 | TTGGGGCT**TGTGT**TCTTTG | 16241 | 16259 |
| 202 | TGGGGCT**TGTGT**TCTTTGC | 16242 | 16260 |
| 203 | GGGCT**TGTGT**TCTTTGCAA | 16244 | 16262 |
| 204 | GGCT**TGTGT**TCTTTGCAAT | 16245 | 16263 |
| 205 | GCT**TGTGT**TCTTTGCAATT | 16246 | 16264 |
| 206 | CT**TGTGT**TCTTTGCAATTC | 16247 | 16265 |
| 207 | CATTAGTTTTCCAT**TGTGT** | 16470 | 16488 |
| 208 | ATTAGTTTTCCAT**TGTGT**G | 16471 | 16489 |
| 209 | TTAGTTTTCCAT**TGTGT**GC | 16472 | 16490 |
| 210 | TAGTTTTCCAT**TGTGT**GCT | 16473 | 16491 |

| | | | |
|---|---|---|---|
| 211 | GTTTTCCAT**TGTGT**GCTAA | 16475 | 16493 |
| 212 | TTTCCAT**TGTGT**GCTAATG | 16477 | 16495 |
| 213 | TCCAT**TGTGT**GCTAATGGA | 16479 | 16497 |
| 214 | **TGTGT**TGGTAGCGATAATG | 16525 | 16543 |
| 215 | TTTCAATTCAG**TGTGT**AGA | 17499 | 17517 |
| 216 | TTCAG**TGTGT**AGACTTATG | 17505 | 17523 |
| 217 | CTGAAGGTTTA**TGTGT**TGA | 18143 | 18161 |
| 218 | GAAGGTTTA**TGTGT**TGACA | 18145 | 18163 |
| 219 | AGGTTTA**TGTGT**TGACATA | 18147 | 18165 |
| 220 | TA**TGTGT**TGACATACCTGG | 18152 | 18170 |
| 221 | AGGTTCATCTAAG**TGTGT**G | 20475 | 20493 |
| 222 | GGTTCATCTAAG**TGTGTGT** | 20476 | 20494 |
| 223 | GTTCATCTAAG**TGTGTGT**G | 20477 | 20495 |
| 224 | TCATCTAAG**TGTGTGTGT**T | 20479 | 20497 |
| 225 | ATCTAAG**TGTGTGTGT**TCT | 20481 | 20499 |
| 226 | CTAAG**TGTGTGTGT**TCTGT | 20483 | 20501 |
| 227 | AAG**TGTGTGTGT**TCTGTTA | 20485 | 20503 |
| 228 | AG**TGTGTGTGT**TCTGTTAT | 20486 | 20504 |
| 229 | G**TGTGTGT**GTTCTGTTATT | 20487 | 20505 |
| 230 | **TGTGTGT**GTTCTGTTATTG | 20488 | 20506 |
| 231 | G**TGTGTGT**TCTGTTATTGA | 20489 | 20507 |
| 232 | **TGTGTGT**TCTGTTATTGAT | 20490 | 20508 |
| 233 | G**TGTGT**TCTGTTATTGATT | 20491 | 20509 |
| 234 | **TGTGT**TCTGTTATTGATTT | 20492 | 20510 |
| 235 | AGTCTCTAGTCAG**TGTGT**T | 21592 | 21610 |
| 236 | TCTCTAGTCAG**TGTGT**TAA | 21594 | 21612 |
| 237 | TCTAGTCAG**TGTGT**TAATC | 21596 | 21614 |
| 238 | TAGTCAG**TGTGT**TAATCTT | 21598 | 21616 |
| 239 | ATCTTAGGGAATT**TGTGT**T | 22125 | 22143 |
| 240 | TCTTAGGGAATT**TGTGT**TT | 22126 | 22144 |
| 241 | CTTAGGGAATT**TGTGT**TTA | 22127 | 22145 |
| 242 | TAGGGAATT**TGTGT**TTAAG | 22129 | 22147 |
| 243 | GGGAATT**TGTGT**TTAAGAA | 22131 | 22149 |
| 244 | ATT**TGTGT**TTAAGAATATT | 22135 | 22153 |
| 245 | TT**TGTGT**TTAAGAATATTG | 22136 | 22154 |
| 246 | T**TGTGT**TTAAGAATATTGA | 22137 | 22155 |
| 247 | **TGTGT**TTAAGAATATTGAT | 22138 | 22156 |
| 248 | C**TGTGT**TGCTGATTATTCT | 22642 | 22660 |
| 249 | **TGTGT**TGCTGATTATTCTG | 22643 | 22661 |
| 250 | ATGTCAGAG**TGTGT**ACTTG | 24647 | 24665 |
| 251 | GTCAGAG**TGTGT**ACTTGGA | 24649 | 24667 |
| 252 | ACATT**TGTGT**CTGGTAACT | 24920 | 24938 |
| 253 | ATT**TGTGT**CTGGTAACTGT | 24922 | 24940 |
| 254 | TT**TGTGT**CTGGTAACTGTG | 24923 | 24941 |
| 255 | T**TGTGT**CTGGTAACTGTGA | 24924 | 24942 |
| 256 | **TGTGT**CTGGTAACTGTGAT | 24925 | 24943 |
| 257 | TGGAGTAAAAGAC**TGTGT**T | 25977 | 25995 |
| 258 | GAGTAAAAGAC**TGTGT**TGT | 25979 | 25997 |
| 259 | GTAAAAGAC**TGTGT**TGTAT | 25981 | 25999 |
| 260 | **TGTGT**TGTATTACACAGTT | 25990 | 26008 |
| 261 | ACTGCGCTTCGAT**TGTGT**G | 26347 | 26365 |
| 262 | CTGCGCTTCGAT**TGTGT**GC | 26348 | 26366 |
| 263 | TGCGCTTCGAT**TGTGT**GCG | 26349 | 26367 |

| | | | |
|---|---|---|---|
| 264 | CGCTTCGAT**TGTGT**GCGTA | 26351 | 26369 |
| 265 | CTTCGAT**TGTGT**GCGTACT | 26353 | 26371 |
| 266 | TCGAT**TGTGT**GCGTACTGC | 26355 | 26373 |
| 267 | GAT**TGTGT**GCGTACTGCTG | 26357 | 26375 |
| 268 | T**TGTGT**GCGTACTGCTGCA | 26359 | 26377 |
| 269 | CCAAGAG**TGTGT**TAGAGGT | 27453 | 27471 |
| 270 | AG**TGTGT**AACATTAGGGAG | 29687 | 29705 |
| 271 | G**TGTGT**AACATTAGGGAGG | 29688 | 29706 |

**Supplemental Table 1**

**List of SARS-CoV-2 GU-rich sequences enriched in Interferon Induction Motif (IIM)**

The reference SARS-CoV-2 genome (NC_045512, positive strand) was scanned for GU-rich ssRNA as explained in Material and Methods. Within the identified 491 GU-rich sequences, 271 were further selected based on their content of at least one "UGUGU" Interferon Induction Motif (IIM). Base numbering is based on the SARS-CoV-2 reference genome NC_045512.
